# Chromosome 2q31.2 Deletion Impairs Neuronal Plasticity through Integrated Stress Response Activation

**DOI:** 10.64898/2026.04.24.720735

**Authors:** Brooke N. Nakamura, Saman Sedighi, Diganta Das, Radhika M. Joshi, Josh Neman

**Affiliations:** Department of Neurological Surgery, USC Brain Tumor Center, Norris Comprehensive Cancer Center, Keck School of Medicine, University of Southern California, Los Angeles, CA, USA

**Keywords:** Keyword: Chromosome 2q31.2 deletion, Integrated stress response, ISRB, ATF4, Synaptic plasticity, Neuronal plasticity, iPSC

## Abstract

Rare genomic deletions disrupt gene networks essential for neural development and synaptic plasticity, yet their mechanistic effects remain poorly understood. Here, we investigated the impact of a chromosome 2q31.2 deletion using induced pluripotent stem cell (iPSC)–derived neural models from an affected individual and a genetically matched parental control.

Differentiation into neural precursor cells and glial lineages was maintained, with subtle changes in cytoskeletal and lineage-associated gene expression. Transcriptomic analysis showed activation of oxidative stress, mitochondrial dysfunction, and degeneration-linked pathways, consistent with engagement of the integrated stress response (ISR). Pharmacologic inhibition of the ISR with ISRIB reduced ATF4 expression in neural precursors and mature neurons and increased CREB phosphorylation throughout development. Functionally, ISRIB improved neuronal network activity in cortical organoids, increasing calcium burst amplitude and synchronization after NMDA and AMPA stimulation. These results identify ISR dysregulation as a mechanistic link between chromosome 2q31.2 deletion and impaired neuronal plasticity, highlighting ISR modulation via ISRIB as a potential therapeutic approach.

## Introduction

Rare genomic disorders offer a powerful framework for uncovering the molecular mechanisms that govern human brain development and function. Structural chromosomal variants, including microdeletions, can disrupt gene networks involved in neuronal differentiation, synaptic signaling, and neural plasticity. However, the mechanistic understanding of many rare deletions remains limited because of small patient populations and the difficulty of modeling early human neurodevelopment in physiologically relevant systems. The chromosomal region 2q31.2 contains genes implicated in transcriptional regulation, cellular stress signaling, and neural development, and deletions within this locus have been associated with neurodevelopmental abnormalities. Despite these associations, the cellular and molecular mechanisms underlying neurological dysfunction in 2q31.2 deletions remain poorly defined. Advances in induced pluripotent stem cell (iPSC) now enable modeling of patient-specific neurodevelopment. Human iPSCs can be differentiated into multiple neural lineages, including neurons and glia, providing a platform to study the functional consequences of genetic variants in a human context (1). Importantly, iPSC-derived neural systems allow interrogation of dynamic signaling pathways that regulate neuronal maturation, synaptic function, and plasticity.

One pathway that has emerged as a critical regulator of neuronal plasticity is the Integrated Stress Response (ISR). The ISR is a conserved adaptive signaling network activated by diverse cellular stressors, including oxidative stress, mitochondrial dysfunction, and metabolic imbalance. Central to this pathway is the phosphorylation of eIF2α, which reduces global protein synthesis while selectively promoting the translation of stress-responsive transcripts such as ATF4 (2,3). While transient ISR activation is protective, chronic activation has been shown to impair synaptic plasticity and memory formation, in part by suppressing CREB-dependent transcriptional programs required for activity-dependent gene expression (4,5).

Pharmacologic inhibition of the ISR with ISRIB (Integrated Stress Response Inhibitor) has been shown to restore translational control downstream of eIF2α phosphorylation and to enhance cognitive function and synaptic plasticity across multiple experimental models (6,7).

These findings suggest that dysregulated ISR signaling may be a key mechanism linking cellular stress to impaired neuronal function.

Based on this framework, we hypothesized that a deletion at 2q31.2 induces neuronal dysfunction through chronic ISR activation, leading to ATF4-mediated suppression of plasticity pathways. To test this hypothesis, we generated iPSC-derived neural models from an individual harboring the deletion (C2074-2qdel) and compared them with a genetically matched parental control (C2075-Paternal). We conducted proteomic pathway analysis, neuronal differentiation assays, protein-level signaling studies, and functional calcium imaging in cortical organoids to determine whether pharmacologic inhibition of the ISR with ISRIB could restore neuronal plasticity.

Our findings demonstrate that ISR signaling is activated in neural cells with a 2q31.2 deletion, resulting in increased ATF4 expression, suppression of CREB-dependent signaling, and impaired activity-dependent neuronal function. Importantly, pharmacologic inhibition of the ISR restores both the molecular and functional features of synaptic plasticity. These results identify ISR dysregulation as a mechanistic link between a rare genomic deletion and impaired neuronal plasticity and highlight ISR modulation as a potential therapeutic strategy for neurodevelopmental disorders characterized by disrupted synaptic function.

## Methods

### iPSC Generation and Characterization

Induced pluripotent stem cells (iPSCs) were generated from peripheral blood mononuclear cells (PBMCs) obtained from the 2q31.2 deletion donor (C2074-2qdel) and a genetically matched parental control (C2075-Paternal) by Applied StemCell, Inc., using a non-integrating Sendai viral reprogramming system. This method enables efficient delivery of pluripotency factors without genomic integration (8). For each donor, the top three iPSC clones were selected based on morphology and growth characteristics. Clones were expanded and cryopreserved (1 × 10 cells per vial; five vials per clone for C2074-2qdel and two vials per clone for C2075-Paternal). Quality control (QC) included pathogen screening (including COVID-19), mycoplasma testing, and recovery assays for each clone. Pluripotency was confirmed by alkaline phosphatase (AP) staining and immunostaining for TRA-1-81 and OCT4. Morphology was assessed using brightfield imaging, and G-banding karyotyping was performed on one representative clone per donor to confirm chromosomal integrity. These validated iPSC lines were used for all downstream neural differentiation and functional analyses.

*Maintenance of Human Pluripotent Stem Cells:* All human iPS cell lines were maintained in mTeSR Plus Medium (STEMCELL Technologies) per the manufacturer’s instructions. Briefly, all cultureware was coated with 1% Geltrex LDEV-Free, hESC-Qualified, Reduced Growth Factor Basement Membrane Matrix (Thermo) in DMEM/F12 + 15 mM HEPES (Thermo) for 1 hour at 37°C before use. Human iPS cell lines were grown in mTeSR medium and passaged using ReLeSR (STEMCELL Technologies) at a ratio of 1:20 to 1:40 every 5-7 days.

*Generation of Neural Progenitor Cells:* To generate neural progenitor cells, we used the STEMdiff SMADi Neural Induction kit with the monolayer culture protocol. All cultureware was coated with 15 ug/mL Poly-L-ornithine (PLO) in PBS for 2 hours at room temperature. PLO was removed, and surfaces were rinsed twice with PBS. The surfaces were then coated with 10 ug/mL laminin in DMEM/F12 with 15 mM HEPES for 2 hours at room temperature. On Day 0, 2×10^6 hiPSC cells were plated in 1 coated well of a 6-well plate in STEMdiff Neural Induction Medium + SMADi + 10 uM Y-27632 and placed in a 37°C 5% CO2 incubator. On Days 1-6, complete media changes were performed daily without 10 uM Y-27632. On Day 7, NPCs were passaged using ACCUTASE. 2×10^6 cells in STEMdiff Neural Induction Medium + SMADi were plated in 1 coated well of a 6-well plate and placed in a 37°C 5% CO2 incubator. On Days 7-13, complete media changes were performed daily. On Day 14, NPCs were passaged using ACCUTASE. 2×10^6 cells in STEMdiff Neural Induction Medium + SMADi were plated in 1 coated well of a 6-well plate and placed in a 37°C 5% CO2 incubator. On Days 15-21, complete media changes were performed daily. On Day 21, NPCs can be used for Forebrain Neuron Differentiation.

*Differentiation of NPCs to Neuronal Forebrain Precursors and Neurons:* To generate neuronal forebrain precursors, we used the STEMdiff Forebrain Neuron Differentiation Kit following the manufacturer’s instructions. All cultureware was coated with 15 ug/mL PLO in PBS for 2 hours at room temperature. PLO was removed, and surfaces were rinsed twice with PBS. The surfaces were then coated with 5 ug/mL laminin in DMEM/F12 with 15 mM HEPES for 2 hours at room temperature. 3.1×10^6 D21 NPCs were plated into a coated T25 flask. Full media changes were performed daily for the next 7 days to differentiate NPCs into neural forebrain precursors. 5.7×10^4 neuronal precursors were plated into each well of a 24-well plate. Full media changes were performed every 2-3 days for at least 8 days to mature neurons.

*ISRIB Treatment*: Forebrain neuronal precursor cells were plated at a density of 57,000 cells per well on poly-L-ornithine/laminin-coated glass coverslips in 24-well plates. Cells were matured for 14 days under forebrain neuronal differentiation conditions to generate mature forebrain neurons. Mature forebrain neurons were treated daily with ISRIB (200 nM; Millipore Sigma, #SML0843) for 7 days. Vehicle-treated cultures were maintained in parallel under identical conditions. Following treatment, coverslips were fixed and processed for qPCR and immunocytochemistry to assess ISR- and synaptic plasticity-associated markers.

*Real-time Quantitative PCR Analysis:* qPCR was carried out to according on a previously established protocol (9–11). Briefly, cells were harvested from culture and RNA was extracted using RNeasy Plus Mini Kit (Qiagen). cDNA synthesis was completed using Maxima First Strand cDNA Synthesis Kit for RT-qPCR (Thermo Scientific). RT-qPCR was then performed using PowerUp SYBR Green Mater Mix (Thermo Scientific) on QuantStudio 6 Flex Real-Time PCR System (Thermo Scientific). All qPCR reactions were run in triplicate, normalized to housekeeping gene expression, and analyzed using the ddCT method. Primer list can be found in Table 1 below.

**Table 1:** Primer List.

| Primer Target Gene: | IDT PrimerTime Assay: |
| --- | --- |
| ACTB | Hs.PT.39a.22214847 |
| CXCR4 | Hs.PT.58.27595676.g |
| FUT4 | Hs.PT.58.14928859.g |
| GAPDH | Hs.PT.39a.22214836 |
| GFAP | Hs.PT.58.1057167 |
| MAP2 | Hs.PT.58.1947612 |
| MSI1 | Hs.PT.58.25123106 |
| NANOG | Hs.PT.58.1185097 |
| NES | Hs.PT.58.1185097 |
| NKX2-2 | Hs.PT.58.25123645.g |
| POU5F1 | Hs.PT.58.14494169.g |
| RELN | Hs.PT.58.39779292 |
| RPLP0 | Hs.PT.39a.22214824 |
| SOX1 | Hs.PT.58.28041414.g |
| SOX2 | Hs.PT.58.237897.g |
| SOX4 | Hs.PT.58.24974948.g |
| SOX9 | Hs.PT.58.38984663 |
| VIM | Hs.PT.58.38906895 |

*Proteomic Analysis:* Proteomic profiling was at ProteasHealth performed using the Biofluid Total Analytic System (BioTAS) platform as previously described (Manousopoulou et al., bioRxiv, 2026; doi: 10.64898/2026.02.16.706239). Briefly, samples were subjected to automated fractionation using a lab-on-chip open tubular chromatography system, followed by reduction, alkylation, and LysC/trypsin digestion. Resulting peptides were labeled with TMTpro isobaric reagents (Thermo Fisher Scientific) for multiplexed quantitative analysis.

Phosphopeptides were enriched using TiO_₂_/ZrO_₂_-based chromatography, and samples were analyzed by ultra-high-performance liquid chromatography (EASY-nLC 1200) coupled to an Exploris 480 Orbitrap mass spectrometer with FAIMS. Raw data were processed using Proteome Discoverer (v2.5) with Sequest search against the UniProt human database, applying a false discovery rate (FDR) of 1% at the peptide and protein levels. Differential protein expression and downstream pathway enrichment analyses were performed to identify biological processes altered in C2074-2qdel relative to C2075-Paternal neural cells.

*Immunocytochemistry staining:* Immunocytochemistry was performed on D21 NPC, D29 FBP, and D42 FBN based on the previously established protocol(12,13). Briefly, cells were cultured on PLO/laminin coated glass coverslips. Cells were fixed with 4% paraformaldehyde for 10 minutes at room temperature and washed twice with PBS. Then cells were permeabilized with 0.3% Triton X-100 in PBS for 30 minutes at 37°C. Cells were then blocked using 1:1 dilution of SeaBlock:PBS for 1 hour at room temperature. Primary antibody was incubated on the cells overnight at 4°C. Secondary antibodies at 1:300 concentration were incubated on the cells for 1 hour at room temperature in the dark. ProLong Gold with DAPI was used to mount the coverslips onto slides. Quantification of immunofluorescent signal was performed as previously described (10,11,14).

*Calcium Imaging in Cortical Organoids:* Induced pluripotent stem cells derived from the 2q31.2 deletion line (C2074-2qdel) and the genetically matched parental control (C2075-Paternal) were differentiated into cortical organoids using a previously established protocol (15) with minor modifications. Briefly, organoid differentiation was initiated with the addition of Noggin (100 ng/mL) during the first 18 days to promote neural induction, followed by long-term culture for 60–120 days to enable neuronal maturation. At day 120, organoids were treated with ISRIB (0.2 µM) or vehicle (DMSO) for 6 hours prior to imaging. For functional calcium imaging, organoids were transduced with pAAV-CAG-SomaGCaMP6f2 (Addgene #158757) to enable visualization of neuronal activity. Following sufficient GFP expression, organoids were transferred to a temperature-controlled recording chamber maintained at 37°C (TC-324C, Warner Instruments) and imaged in BrainPhys Imaging Optimized Medium (STEMCELL Technologies, #05796). Neuronal activity was evoked using NMDA (100 µM) or AMPA (100 µM) applied via bath perfusion. Imaging was performed using a Leica SP-8X multiphoton microscope equipped with a 25× 0.95 NA water-immersion objective (2.5 mm working distance), capturing a field of view of approximately 200 × 200 µm². Time-lapse recordings were acquired at ∼1 frame per 860 ms. Calcium imaging data (raw TIFF format) were analyzed in MATLAB (MathWorks) using the CaImAn toolbox, which applies constrained non-negative matrix factorization (CNMF) to extract regions of interest (ROIs), identify calcium transients, and infer spike activity (16).

Calcium traces were normalized and expressed as relative fluorescence changes over time. Organoids were derived from a single differentiation batch, including three baseline organoids, three NMDA-stimulated organoids, and one AMPA-stimulated organoid. For each organoid, three independent imaging recordings were acquired to account for intra-organoid variability due to spatial sampling limitations. Thus, analyses reflect technical replicates within a single differentiation batch.

*Statistical Analysis*: All data are presented as mean ± standard error of the mean (SEM) unless otherwise indicated. Statistical analyses were performed using GraphPad Prism. For comparisons between two groups, two-tailed unpaired Student’s t-tests were used when data were normally distributed. For comparisons involving more than two groups or multiple conditions (e.g., differentiation stages or treatment groups), one-way or two-way analysis of variance (ANOVA) was performed, followed by Tukey’s or Bonferroni post hoc tests to correct for multiple comparisons. All statistical significance was defined as: *p < 0.05, **p < 0.01, ***p < 0.001, ****p < 0.0001.

Proteomic data were analyzed using standard quantitative workflows for TMTpro-labeled mass spectrometry datasets. Data were first log2-transformed and normalized to correct for loading and labeling efficiency across samples. Differential protein expression between C2074-2qdel and C2075-Paternal samples was assessed using two-tailed Student’s t-tests or linear models, where appropriate. Multiple hypothesis testing was controlled using the Benjamini–Hochberg false discovery rate (FDR) correction, with a threshold of FDR < 0.05 considered statistically significant. For pathway enrichment analysis, significantly altered proteins were mapped to canonical pathways using curated databases (e.g., Ingenuity Pathway Analysis or equivalent). Enrichment significance was calculated using right-tailed Fisher’s exact test, and results are reported as −log(p-value). A threshold of −log(p-value) ≥ 1.3 (p ≤ 0.05) was used to define significantly enriched pathways. Only proteins identified with ≥1 unique peptide and meeting quality filters (peptide and protein FDR ≤ 1%) were included in downstream analyses. Data are presented as mean ± SD unless otherwise stated.

Calcium imaging data were analyzed at the level of individual regions of interest (ROIs) and aggregated per organoid. Comparisons of event amplitude, burst frequency, and synchrony between ISRIB-treated and control groups were performed using unpaired t-tests or Mann–Whitney U tests when normality assumptions were not met. Normality of datasets was assessed using the Shapiro–Wilk test, and variance equality was evaluated using Levene’s test.

## Results

### Generation and Characterization of Patient-Derived Neural Lineages Harboring the Chromosome 2q31.2 Deletion

To investigate the cellular consequences of chromosome 2q31.2 deletion on neural development, induced pluripotent stem cells (iPSCs) were created from an affected individual (C2074-2qdel) and a genetically related parental control (C2075-Paternal). To assess transcriptional changes during neural differentiation, quantitative PCR (qPCR) was performed comparing iPSCs and day 21 neural precursor cells (NPCs) derived from both lines (Figure 1). Pluripotency-related genes POU5F1 (OCT4), FUT4, and NANOG were highly expressed in iPSCs and significantly decreased upon differentiation into NPCs (Figure 1A). Early neural transcription factors SOX1, SOX2, SOX4, SOX9, MSI1, NKX2-2, and NOTCH1 were strongly upregulated in NPCs compared to iPSCs in both groups (Figure 1B). Additional mRNA analysis of neural precursor identity revealed significant increases in cytoskeletal and lineage-specific Vimentin and Nestin (Figure 1C), confirming NPC identity. Genes involved in neuronal migration and cortical organization, including CXCR4 and RELN, were significantly upregulated in C2074-2qdel showing higher expression than C2075-Paternal. MAP2, a marker of early neuronal maturity, was detectable at this point, indicating progression toward neuronal differentiation.

**Figure 1.**
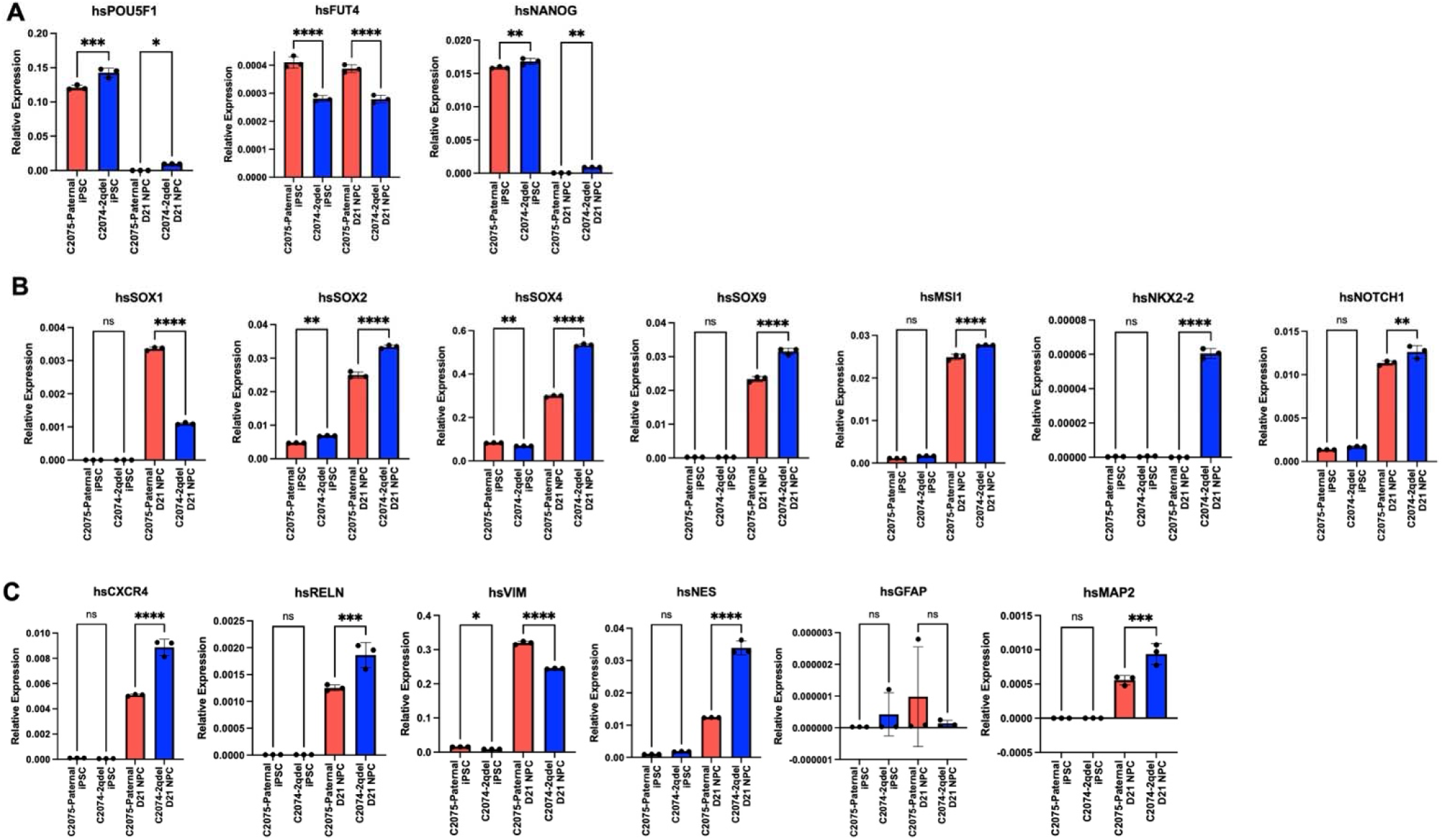
qPCR analysis of neural differentiation from iPSCs harboring chromosome 2q31.2 deletion. Quantitative PCR (qPCR) analysis of gene expression during differentiation of iPSCs from the 2q31.2 deletion line (C2074-2qdel) and a genetically matched parental control (C2075-Paternal) into day 21 neural precursor cells (NPCs). Data are presented as mean ± SEM. (A) Pluripotency markers (POU5F1, FUT4, NANOG) are highly expressed in iPSCs and significantly downregulated upon differentiation into NPCs in both control and patient lines, confirming loss of pluripotency. (B) Neural transcription factors (SOX1, SOX2, SOX4, SOX9, MSI1, NKX2-2, NOTCH1) are robustly upregulated in NPCs, indicating successful neural lineage specification in both groups. (C) Neural precursor and cytoskeletal markers (CXCR4, RELN, VIM, NES, GFAP, MAP2) are increased following differentiation, with patient-derived NPCs showing gene-specific differences, particularly in migration and cytoskeletal-associated genes. Statistical significance was determined by two-tailed t-test or ANOVA (*p < 0.05, **p < 0.01, ***p < 0.001, ****p < 0.0001; ns = not significant).

Conversely, GFAP expression remained low, consistent with limited astrocyte differentiation in early NPC cultures. Overall, these findings reveal that neural differentiation is efficient in 2q31.2-deletion cells, but transcriptional differences in cytoskeletal and migration pathways suggest early changes in neural developmental programs.

To determine whether these transcriptional differences affect lineage potential, immunofluorescence analysis was performed to assess differentiation into various central nervous system lineages (Figure 2). NPCs derived from both C2074-2qdel and C2075-Paternal showed strong Nestin and GFAP expression, confirming neural precursor identity (Figure 2A).

**Figure 2.**
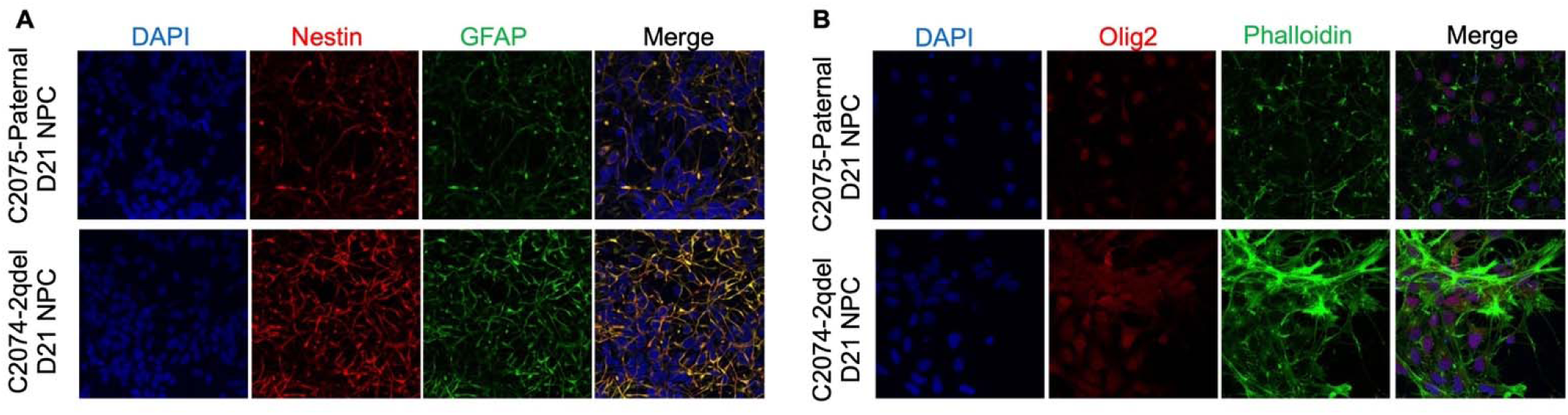
Immunofluorescence characterization of iPSC-derived neural precursor cells from 2q31.2 deletion and control lines. Immunofluorescence analysis of day 21 neural precursor cells (NPCs) derived from the 2q31.2 deletion line (C2074-2qdel) and the genetically matched parental control (C2075-Paternal). (A) NPC identity and astrocytic lineage markers. Cells were stained for Nestin (neural precursor marker) and GFAP (astrocytic marker), with DAPI labeling nuclei. Both lines exhibit robust Nestin and GFAP expression, confirming neural precursor identity. (B) Oligodendrocyte lineage and cytoskeletal organization. Cells were stained for Olig2 (oligodendrocyte lineage marker) and phalloidin (F-actin cytoskeleton), with DAPI labeling nuclei. Both lines show Olig2-positive cells, confirming oligodendrocyte lineage potential.

The oligodendrocyte marker Olig2 further confirmed the ability of both C2074-2qdel and C2075-Paternal NPCs to produce oligodendroglial progenitors (Figure 2B). Olig2-positive cells were evenly spread throughout the cultures, showing efficient lineage specification. Co-staining with phalloidin revealed well-structured actin cytoskeletons in both groups; however, patient cells exhibited more prominent cytoskeletal organization, with more cellular projections and more complex networks. These structural differences imply that the chromosome 2q31.2 deletion may impact cytoskeletal dynamics and cell morphology during neural development.

Overall, these results show that iPSCs with a deletion at 2q31.2 can still differentiate into neural precursor cells and generate multiple CNS lineages, including astrocytes and oligodendrocytes. However, subtle yet consistent differences in transcriptional profiles, lineage marker expression, and cytoskeletal organization indicate that the deletion influences early neural development, potentially leading to downstream issues in neuronal maturation and synaptic plasticity.

### Proteomic Analysis Identifies Activation of Integrated Stress Response and Degeneration-Associated Pathways in 2q31.2 Deletion Neural Cells

To determine how the 2q31.2 deletion alters molecular programs during early neural development, we performed protein-level pathway enrichment analysis of day 21 neural precursor cells derived from C2074-2qdel and C2075-Paternal. Differential protein expression and downstream pathway analysis revealed a clear, coordinated shift toward activation of cellular stress, metabolic dysfunction, and degeneration-associated signaling pathways in C2074-2qdel cells (Figure 3). Across all pathways, enrichment exceeded the statistical threshold (−log(p-value) > ∼1.3), with multiple pathways showing strong significance (−log(p-value) > 3–5), indicating robust, reproducible protein-level alterations.

**Figure 3.**
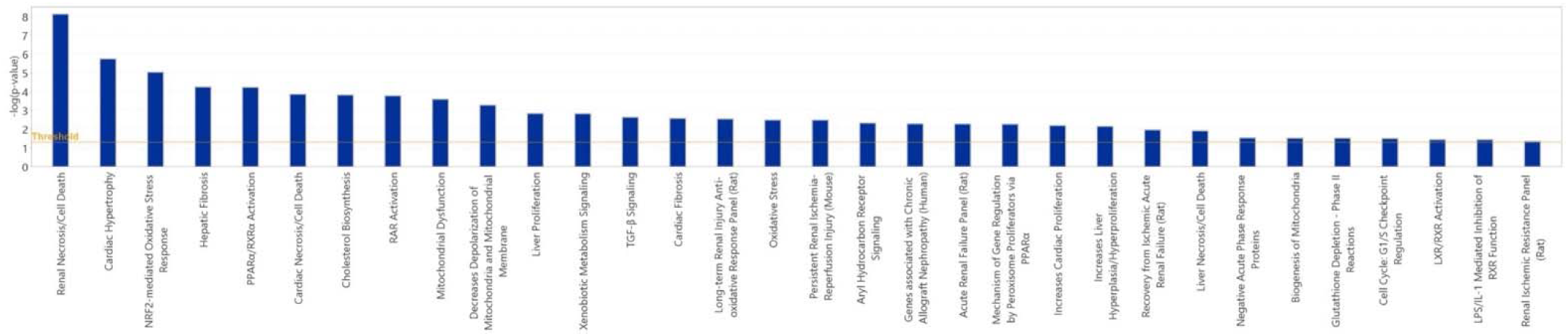
Proteomic pathway enrichment reveals activation of stress-and degeneration-associated signaling in neural cells with a 2q31.2 deletion. Proteomic pathway enrichment analysis comparing day 21 neural precursor cells derived from the 2q31.2 deletion line (C2074-2qdel) and the genetically matched parental control (C2075-Paternal). Differential protein expression was analyzed and mapped to canonical pathways, with significance represented as−log(p-value). Top enriched pathways in C2074-2qdel cells include NRF2-mediated oxidative stress response, necrosis/cell death signaling, mitochondrial dysfunction, and metabolic and nuclear receptor signaling pathways (e.g., PPAR/RXR and RAR signaling). Additional enrichment of pathways associated with cellular injury, inflammation, and tissue remodeling is observed, reflecting activation of conserved stress-response programs. The horizontal threshold line indicates the cutoff for statistical significance (−log(p-value) ≈ 1.3).

Among the most significantly enriched pathways, the NRF2-mediated oxidative stress response was the most altered, indicating elevated oxidative stress signaling in NPCs with a 2q31.2 deletion. This was accompanied by enrichment of pathways associated with cellular injury and death, including necrosis/cell death signaling and broader oxidative damage networks, suggesting that patient-derived neural cells exist in a heightened basal stress state even at early stages of differentiation. In parallel, strong enrichment of mitochondrial dysfunction, mitochondrial membrane depolarization, and xenobiotic metabolism signaling indicates disruption of cellular energetics and metabolic homeostasis.

Importantly, these proteomic signatures converge on biological processes known to activate the integrated stress response (ISR). Oxidative stress and mitochondrial dysfunction are well-established triggers of eIF2α phosphorylation, which attenuates global protein synthesis while selectively promoting the translation of stress-adaptive transcripts, including the transcription factor ATF4 (3). Consistent with this, enrichment of mitochondrial and metabolic stress pathways suggests upstream activation of eIF2α kinases and engagement of ISR signaling. ATF4, a central ISR effector, regulates transcriptional programs that prioritize stress adaptation but suppress CREB-dependent plasticity pathways when chronically activated(4).

Additional dysregulated pathways—including PPARα/RXRα activation, RAR signaling, cholesterol biosynthesis, and LPS/IL-1–mediated inhibition of RXR function—further support a cellular environment marked by metabolic reprogramming, inflammatory signaling, and chronic stress adaptation. The enrichment of pathways typically associated with tissue injury and remodeling, such as hepatic fibrosis and cardiac hypertrophy, likely reflects activation of conserved stress-response programs rather than tissue-specific effects, reinforcing a global shift toward a damage-response phenotype. Notably, pathways related to mitochondrial biogenesis, cellular proliferation, and cytoskeletal organization were also altered, suggesting that stress signaling may affect both structural and developmental processes in neural precursor cells.

Collectively, these data support a model in which a deletion at 2q31.2 induces chronic ISR activation, characterized by eIF2α-mediated translational reprogramming and increased ATF4 signaling, leading to disruption of metabolic homeostasis, mitochondrial function, and neuronal maintenance pathways.

### Stage-Dependent Suppression of ATF4 by ISRIB in 2q31.2 Deletion Neural Cells

Based on proteomic pathway evidence implicating activation of cellular stress programs consistent with engagement of the integrated stress response, we next directly interrogated the ISR effector ATF4 to determine whether pharmacologic modulation of ISR signaling could restore downstream pathways relevant to neuronal plasticity. Neural cultures derived from the 2q31.2 deletion line (C2074-2qdel) and the genetically matched control (C2075-Paternal) were treated with ISRIB, a small molecule that restores translational control downstream of eIF2α phosphorylation.

Immunofluorescence analysis revealed strong nuclear ATF4 expression in C2074-2qdel neural cells across differentiation stages (Figure 4). On day 21, at the neural progenitor stage, both C2074-2qdel and C2075-Paternal cells exhibited strong ATF4 staining; however, ISRIB treatment did not significantly reduce ATF4 levels in C2074-2qdel NPCs (Figure 4A).

**Figure 4.**
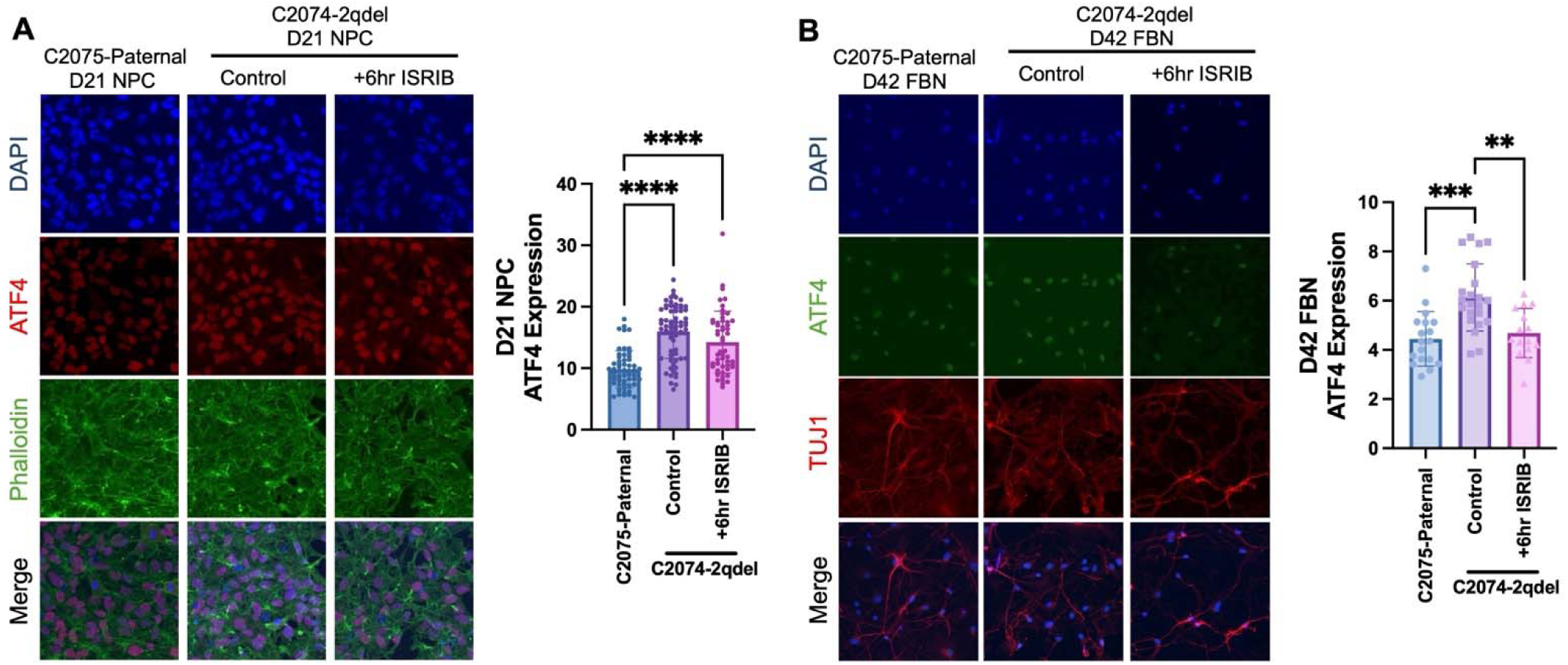
ISRIB induces a developmental-stage-dependent reduction in ATF4 in neural cells with a 2q31.2 deletion. Immunofluorescence analysis of ATF4 expression in neural cells derived from the 2q31.2 deletion line (C2074-2qdel) and the genetically matched parental control (C2075-Paternal) following 6-hour ISRIB treatment. (A) Day 21 neural progenitor cells (NPCs). Cells were stained for ATF4 (red), phalloidin (green, cytoskeleton), and DAPI (blue, nuclei). B) Day 42 forebrain neurons (FBNs). Cells were stained for ATF4 (green), TuJ1 (red, neuronal marker), and DAPI (blue). Quantification of ATF4 intensity is shown on the right. (**p < 0.01, ***p < 0.001, ****p < 0.0001).

Quantitative analysis confirmed no meaningful decrease in ATF4 signal intensity after ISRIB exposure, suggesting that early neural progenitors are relatively refractory to short-term ISR inhibition. This indicates that ATF4 expression at this stage may be maintained by mechanisms that are less dependent on sustained ISR signaling or less sensitive to ISRIB-mediated translational control.

On day 42, ISRIB treatment significantly reduced ATF4 expression in forebrain neurons (FBNs) from C2074-2qdel cells (Figure 4B). ATF4 staining was significantly reduced after ISRIB exposure, particularly in TuJ1-positive neuronal populations, indicating effective suppression of ISR signaling in differentiated neurons. These findings demonstrate that ISR signaling becomes increasingly dependent on eIF2α-mediated translational control during neuronal maturation, making ATF4 expression more sensitive to ISRIB.

Across developmental stages, these results show a progressive increase in ISRIB sensitivity, with minimal effects in early progenitors and pronounced suppression of ATF4 in mature neurons. Mechanistically, this supports a model in which chronic ISR activation—driven by upstream stress signals identified in proteomic analyses—leads to sustained ATF4 expression in 2q31.2 deletion cells, particularly in differentiated neuronal populations. Given that ATF4 is a key negative regulator of synaptic plasticity and suppresses CREB-dependent transcriptional programs, ISRIB-mediated reduction of ATF4 is a critical mechanistic step toward restoring neuronal function.

Together, these findings establish a direct mechanistic link among chromosome 2q31.2 deletion, stress-induced ISR activation, and downstream neuronal dysfunction, and demonstrate that pharmacologic inhibition of the ISR can selectively suppress ATF4 in mature neurons. This stage-dependent modulation of ISR signaling provides a strong rationale for targeting the ISR to restore downstream plasticity pathways in patient-derived neural models.

### ISRIB Enhances CREB Activation Across Neural Differentiation Stages

To determine whether pharmacologic inhibition of the integrated stress response restores downstream plasticity signaling, we assessed CREB activation by measuring phosphorylated CREB (pCREB) levels in neural cells derived from the 2q31.2 deletion line (C2074-2qdel) and the genetically matched control (C2075-Paternal) after ISRIB treatment (Figure 5). CREB phosphorylation is a central regulator of activity-dependent gene expression and synaptic plasticity. At baseline, C2074-2qdel cells showed reduced pCREB levels compared with C2075-Paternal controls, consistent with impaired activation of plasticity-associated signaling pathways.

**Figure 5.**
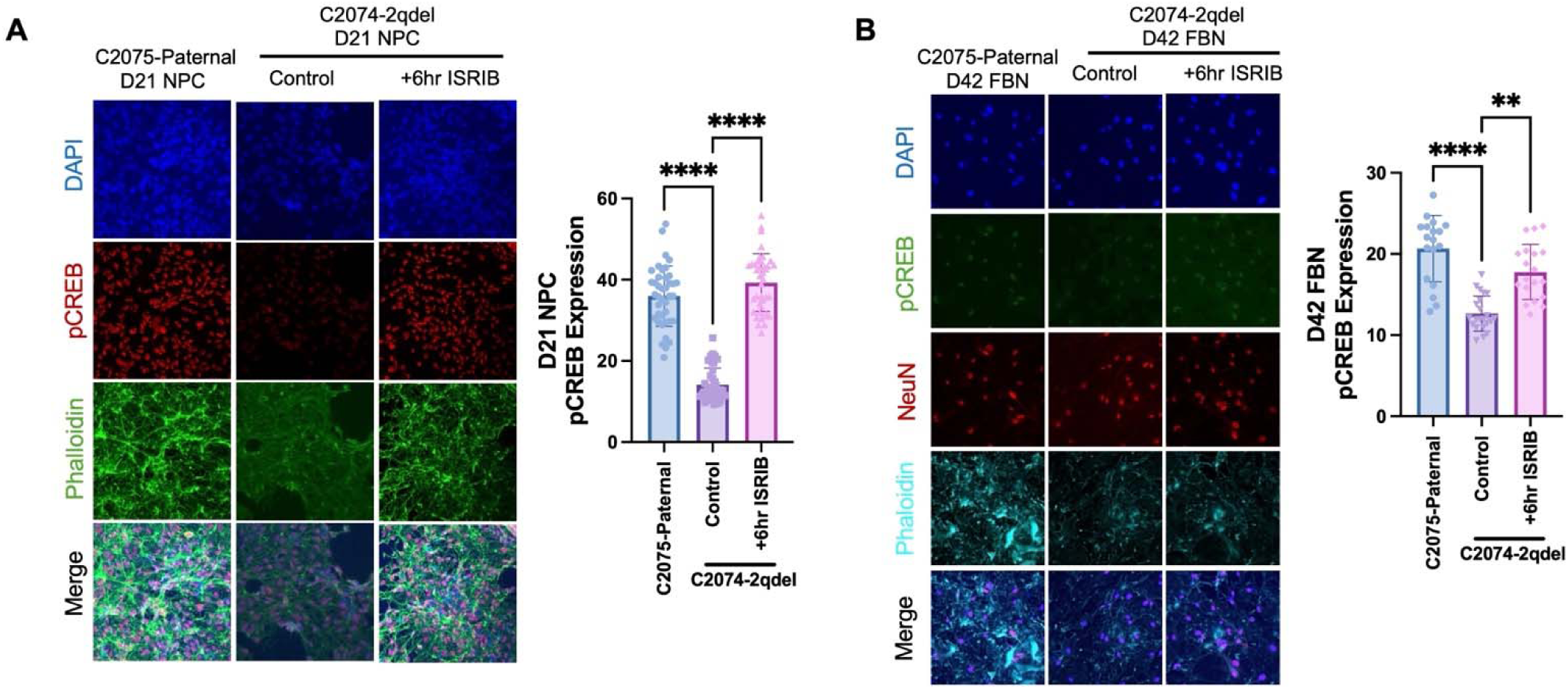
ISRIB enhances CREB activation across neural differentiation stages. Immunofluorescence analysis of phosphorylated CREB (pCREB) in neural cells derived from the 2q31.2 deletion line (C2074-2qdel) and the genetically matched parental control (C2075-Paternal) following 6-hour ISRIB treatment. (A) Day 21 neural progenitor cells (NPCs) show increased pCREB expression following ISRIB treatment. (C) Day 42 forebrain neurons (FBNs) display robust induction of pCREB in NeuN-positive neurons following ISRIB treatment. DAPI marks nuclei and phalloidin labels cytoskeletal structure. Quantification of pCREB intensity is shown on the right. (**p < 0.01, ***p < 0.001, ****p < 0.0001).

This deficit persisted across differentiation stages, including day 21 NPCs (Figure 5A) and day 42 forebrain neurons (Figure 5B). After 6-hour ISRIB treatment, a significant increase in pCREB was observed at both stages.

Notably, increased pCREB was observed in day 21 NPCs, despite the absence of a significant ATF4 reduction at this stage (Figure 4). This suggests that ISRIB-mediated enhancement of CREB signaling occurs through both ATF4-dependent mechanisms—via relief of ISR-mediated translational repression—and ATF4-independent pathways. This is consistent with ISRIB’s broader role in restoring global protein synthesis downstream of eIF2α phosphorylation, thereby relieving repression of plasticity-associated transcripts.

Collectively, these findings demonstrate that ISRIB reverses stress-induced suppression of CREB signaling in neural cells with a 2q31.2 deletion, thereby restoring a key molecular pathway required for synaptic plasticity and neuronal function. These data establish a direct mechanistic link between ISR activation and impaired CREB-dependent plasticity and show that pharmacologic inhibition of the ISR rescues activity-dependent transcriptional programs essential for neuronal signaling.

### ISRIB Enhances Activity-Dependent Neuronal Network Function in Cortical Organoids

To determine whether ISRIB-mediated molecular changes translate into functional improvements in neuronal activity, we performed calcium imaging in 120-day cortical organoids derived from the 2q31.2 deletion line (C2074-2qdel) and the genetically matched control (C2075-Paternal) after pretreatment with ISRIB (0.2 µM, 6 hours) or vehicle (Figure 6).

**Figure 6.**
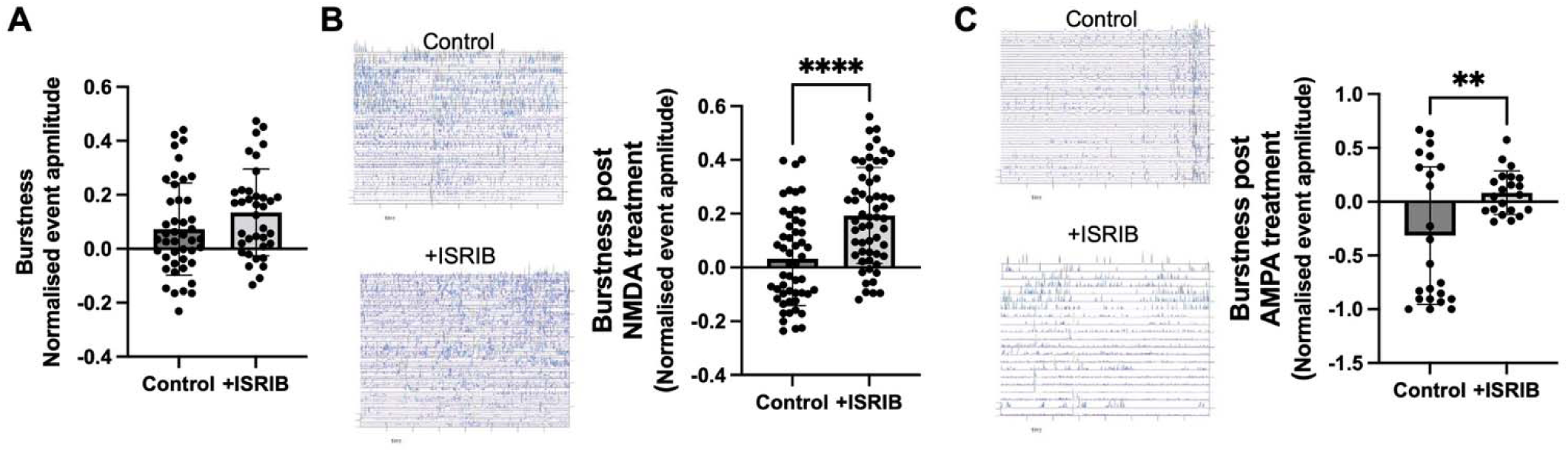
ISRIB enhances activity-dependent neuronal network responses in cortical organoids. Calcium imaging analysis of neuronal activity in 120-day cortical organoids derived from the 2q31.2 deletion line (C2074-2qdel) following treatment with ISRIB (0.2 µM, 6 hours) or vehicle control. (A) Quantification of normalized calcium event amplitude (burstiness) shows no significant difference between control and ISRIB-treated organoids under unstimulated conditions, indicating that ISRIB does not alter spontaneous neuronal activity. (B) Representative calcium traces (left) and quantification (right) demonstrate a significant increase in burst amplitude and network activity in ISRIB-treated organoids compared to control following NMDA (100 µM) stimulation. (C) Representative calcium traces (left) and quantification (right) show enhanced burst amplitude and synchronization in ISRIB-treated organoids following AMPA (100 µM) stimulation. (**p < 0.01, ****p < 0.0001).

At baseline, before glutamatergic stimulation, both ISRIB- and vehicle-treated organoids exhibited spontaneous calcium transients localized to small clusters of neurons, consistent with the presence of intrinsically active microcircuits. Qualitative spatial mapping of active regions of interest (ROIs) revealed no statistically significant differences in baseline activity between treatment groups, indicating that ISR inhibition does not induce nonspecific hyperexcitability or broadly alter spontaneous neuronal firing (Figure 6A; Supplemental Video 1, Supplemental Video2).

In contrast, bath application of NMDA (100 µM) or AMPA (100 µM) elicited a markedly enhanced calcium response in ISRIB-pretreated C2074-2qdel organoids compared with vehicle-treated controls. Following NMDA stimulation (Figure 6B), ISRIB-treated organoids showed increased burst amplitude and network synchronization, with more frequent and coordinated calcium transients across neuronal populations. Quantitative analysis confirmed a significant increase in normalized event amplitude, indicating enhanced excitatory responsiveness.

Similarly, after AMPA stimulation (Figure 6C), ISRIB-treated organoids displayed greater firing intensity and improved temporal coordination of calcium signals compared with untreated controls, with a significant increase in normalized burst amplitude. In contrast, vehicle-treated organoids exhibited lower-amplitude, less synchronized activity.

Across both stimulation paradigms, ISRIB treatment consistently enhanced the magnitude and coordination of stimulus-evoked neuronal activity, indicating improved synaptic integration and excitatory network dynamics. Notably, these functional effects were observed under excitatory challenge rather than at baseline, suggesting that the functional consequences of the 2q31.2 deletion are most apparent during activity-dependent signaling.

These findings are consistent with ISRIB-mediated molecular changes, including reduced ATF4 (Figure 4) and restored CREB phosphorylation (Figure 5), and support a model in which ISR activation suppresses neuronal plasticity via eIF2α-dependent translational repression.

ISRIB, by restoring translational control downstream of eIF2α phosphorylation, selectively enhances stimulus-evoked neuronal responsiveness without altering baseline excitability.

Collectively, these data demonstrate that pharmacologic inhibition of the ISR rescues activity-dependent neuronal network function in cortical organoids derived from cells with a 2q31.2 deletion, providing functional evidence that ISR dysregulation contributes to impaired synaptic plasticity and is therapeutically targetable.

## Discussion

This study identifies activation of the integrated stress response as a key mechanism linking chromosome 2q31.2 deletion to impaired neuronal plasticity. Rather than broadly disrupting neural differentiation, the deletion appears to induce a stress-adapted cellular state that selectively constrains activity-dependent neuronal function.

Proteomic pathway analysis revealed enrichment of oxidative stress, mitochondrial dysfunction, and metabolic dysregulation, indicating that C2074-2qdel neural cells exist in a chronically stressed state. These upstream signals are well-established triggers of eIF2α phosphorylation, leading to activation of the ISR and preferential translation of ATF4 (17).

Consistent with this model, we observed sustained ATF4 expression in patient-derived neural cells, particularly in differentiated neurons. Functionally, this ISR activation appears to repress neuronal plasticity signaling. ATF4 has been shown to suppress CREB-dependent transcription, which is essential for synaptic strengthening, memory-related gene expression, and activity-dependent neuronal adaptation (4,18). Accordingly, C2074-2qdel cells exhibited reduced baseline pCREB, indicating impaired activation of plasticity-associated pathways.

Pharmacologic inhibition of the ISR with ISRIB restored this balance. ISRIB reduced ATF4 levels in mature neurons and robustly increased pCREB across developmental stages, consistent with restoration of translational control and activity-dependent gene expression. Importantly, these molecular effects translated into functional rescue in cortical organoids.

From initial calcium imaging analysis, at baseline, prior to glutamatergic stimulation, both ISRIB- and vehicle-treated organoids exhibited spontaneous calcium transients localized to small clusters of neurons, consistent with the presence of intrinsically active microcircuits.

Quantitative analysis revealed no statistically significant differences in baseline activity between treatment groups, indicating that ISR inhibition does not induce nonspecific hyperexcitability or broadly alter spontaneous neuronal firing. Notably, however, preliminary qualitative assessment suggested that ISRIB-treated organoids exhibited more organized and elongated neurite/axonal structures, consistent with a healthier neuronal network architecture. While these observations require further systematic and quantitative validation, they are consistent with the increased pCREB signaling and restoration of synaptic plasticity pathways observed in treated cells, supporting the interpretation that ISRIB may enhance neuronal health and connectivity even in the absence of measurable changes in baseline activity.

The effect of ISRIB became most apparent under excitatory challenge. Following NMDA and AMPA receptor stimulation, ISRIB-treated organoids showed increased burst amplitude and enhanced network synchronization, suggesting improved glutamatergic responsiveness and activity-dependent network engagement. NMDA and AMPA receptors are central mediators of excitatory synaptic transmission and synaptic plasticity, including long-term potentiation (19,20) Thus, the enhanced responses to NMDA and AMPA stimulation indicate that ISRIB does not simply increase basal excitability, but rather restores the capacity of neuronal networks to respond to synaptic demand.This distinction is critical-Our data indicate that chromosome 2q31.2 deletion does not primarily impair intrinsic neuronal excitability at rest; instead, it limits the dynamic range of neuronal responses during activity-dependent signaling. ISR activation may therefore function as a molecular “brake” on plasticity, restricting the ability of neurons to engage in coordinated network activity. ISRIB appears to release this constraint, restoring a latent functional reserve that becomes apparent during glutamatergic stimulation.

These findings suggest that ISR dysregulation may represent a convergent mechanism underlying neural dysfunction, particularly in conditions where cellular stress intersects with synaptic plasticity. Moreover, proteomic signatures implicating lipid metabolism, mitochondrial pathways, and cytoskeletal remodeling suggest that ISR activation may also affect oligodendrocyte function and white matter integrity. This is an important direction for future work in patients with chromosome 2q31.2 deletion, as chronic ISR activation has been linked to myelin disorders and impaired glial homeostasis (21,22).

Overall, this work supports a model in which chronic ISR activation suppresses CREB-dependent plasticity and limits activity-dependent neuronal function. Targeting translational control pathways with ISRIB restores molecular signaling and network-level responsiveness in this rare genomic disorder model.

## Acknowledgments

The authors gratefully acknowledge the generous support of the Bidsal Family Fund, whose contribution made this study possible. We thank Giorgia Quadrato, Marcella Birtele, and Tuan Nguyen for their valuable contributions to organoid and calcium imaging experiments.

